# Multiscale and extended retrieval of associative memory structures in a cortical model of local-global inhibition balance

**DOI:** 10.1101/2021.11.30.470555

**Authors:** Thomas F Burns, Tatsuya Haga, Tomoki Fukai

**Affiliations:** Neural Coding and Brain Computing Unit, OIST Graduate University, Okinawa, Japan

## Abstract

Inhibitory neurons take on many forms and functions. How this diversity contributes to memory function is not completely known. Previous formal studies indicate inhibition differentiated by local and global connectivity in associative memory networks functions to rescale the level of retrieval of excitatory assemblies. However, such studies lack biological details such as a distinction between types of neurons (excitatory and inhibitory), unrealistic connection schemas, and non-sparse assemblies. In this study, we present a rate-based cortical model where neurons are distinguished (as excitatory, local inhibitory, or global inhibitory), connected more realistically, and where memory items correspond to sparse excitatory assemblies. We use this model to study how local-global inhibition balance can alter memory retrieval in associative memory structures, including naturalistic and artificial structures. Experimental studies have reported inhibitory neurons and their sub-types uniquely respond to specific stimuli and can form sophisticated, joint excitatory-inhibitory assemblies. Our model suggests such joint assemblies, as well as a distribution and rebalancing of overall inhibition between two inhibitory sub-populations – one connected to excitatory assemblies locally and the other connected globally – can quadruple the range of retrieval across related memories. We identify a possible functional role for local-global inhibitory balance to, in the context of choice or preference of relationships, permit and maintain a broader range of memory items when local inhibition is dominant and conversely consolidate and strengthen a smaller range of memory items when global inhibition is dominant. This model therefore highlights a biologically-plausible and behaviourally-useful function of inhibitory diversity in memory.

## Introduction

The mechanisms by which our brains flexibly perform the complex tasks of learning and memory are not completely understood. Hebbian learning [Hebb, 1945], the relative increase in synaptic strength between neurons as a result of shared, causal activity, seems important. Hebb postulated memories were formulated in the brain by assemblies of highly-interconnected neurons [Hebb, 1945]. Evidence for this ‘neuron assembly’ hypothesis was found in hippocampus, where groups of neurons become synchronously activated in response to an animal’s spatial location, indicating a neural correspondence to and potential memory of the location [Harris et al., 2003]. These memories are often mutually related – in physical or behavioural space for the case of navigation [Tolman, 1948], in reward space for the case of rewarded learning tasks [Dusek & Eichenbaum, 1997], in linguistic space for the case of language comprehension [Goldstein et al., 2021], and theoretically in any arbitrary semantic space for generalized graph-based reasoning (e.g., family trees) [Whittington et al., 2020]. How can the structure of these mutual relations be identified dynamically in cortical networks? Inhibitory mechanisms may hold an answer. Here, we computationally explore the possible role of inhibitory circuits in extracting graph-based relationships in the space of behaviorally relevant information.

The majority of experimental and computational work focusing on assemblies as representations of memory items has focused on the role of excitatory neurons. However, emerging evidence suggests inhibitory neurons play a non-trivial role in cortical networks. Throughout the brain, inhibitory neurons have classically been thought to coarsely keep excitation in check with a broad, non-specific *blanket of inhibition* [Brunel, 2000; Amit et al. 1994]. But more recent work has shown inhibitory neurons are tuned to specific external stimuli [Okun & Lampl, 2008; Xue et al., 2014], have specific associations with behaviour [Dudok et al., 2021], have a large diversity of forms and functions within and across brain areas [Gouwens & Sorensen, 2020; Burns & Rajan, 2021], and form inhibitory assemblies [Zhang, 2017], often jointly with excitatory subnetworks [Otsuka & Kawaguchi, 2009; Koolschijn et al., 2019]. A hallmark of many neuropathologies is inhibitory disfunction [Amieva et al., 2004; Baroncelli et al., 2011]. If specific inhibitory disfunction alone sufficient for explaining these pathologies, then we could expect subtle inhibitory changes to cause dramatic changes in global function in complex tasks like those involving learning and memory. A greater understanding of the neurophysiological mechanisms underlying these changes may help us target treatments for such disorders and provide fundamental insight into the computational roles of inhibitory neurons in such circuits.

Previous modelling work in a formal model with binary neurons [Haga & Fukai, 2019] has shown how anti-Hebbian learning (i.e., involving inhibitory synapses) in an associative memory model was able to extend the span of association between mutually-related memory items organised in a simple ring structure, compared to a regular Hebbian learning rule (i.e., not involving inhibitory synapses). Later work extended this formal model to arbitrary graph structures [Haga & Fukai, 2021]. These results suggest inhibition may play a nontrivial role in relational memory systems. However, these models lacked biological features – most prominently a lack of distinction between excitatory and inhibitory neuron populations, breaking Dale’s Law. The excitatory assemblies were also not nearly as sparse as those seen in biology and the neurons took on binary states. Nevertheless, the results indicate global functional changes can result from subtle inhibitory changes. This study proposes a realistic connection scheme of distinct excitatory and inhibitory neurons to embed sparse cell assemblies which represent memory items mutually linked through arbitrary graph structures. Formulated in this way, the model allows us to confirm the previous suggestion that a balance between local inhibition and global inhibition on cell assemblies determines the scale and extent of memories retrieved in a neural network. We show this for various naturalistic and artificial associative memory structures, including as a behaviourally-useful function to maintain a choice distribution given a juncture or decision point in physical or memory space. We find a balance between local and global inhibition allows control over the range of recall within arbitrary graph structures, as well as graph clustering effects which may be useful in navigation and memory tasks.

## Materials and Methods

### Model

In order to embed memories in the network, we generate binary patterns as vectors of length *N*^*E*^, the number of excitatory neurons. Then, the weight *T*_*ij*_ of connections between any pair of excitatory neurons *i* and *j* is defined using these patterns. First, we create *p* random binary patterns (of 0s and 1s) of length 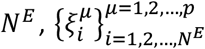, with probabilities for 0 and 1 as 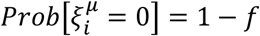 and 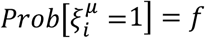, and where we call *f* the “sparseness” parameter of the memory patterns. Memories are then embedded using a modified extended association rule [18, 29] designed to allow association between memory items in an arbitrary graph structure where vertices are the memory patterns and edges represent an association of two memory patterns. Specifically

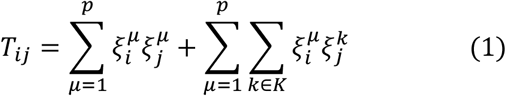

where *K* is the set of memory patterns neighbouring pattern *μ* in the associative memory structure, *M*.

Two populations of inhibitory neurons are also modelled – one with global connectivity (uniform connection probabilities as indicated in Figure 1) of size *N*^*G*^ and another with local connectivity, which is specific to each memory pattern, and has a total size of *N*^*L*^, but where only *fN*^*L*^ local inhibitory neurons participate in each pattern. Unless stated otherwise, we use *N*^*E*^ = 4,000, *N*^*G*^ = 500, *N*^*L*^ = 500, and *f* = 0.01, meaning that each pattern consists of a joint assembly of 40 excitatory neurons and 10 local inhibitory neurons. A general schematic of the model from the perspective of a single memory pattern is shown in Figure 1A.

**Figure 1.**
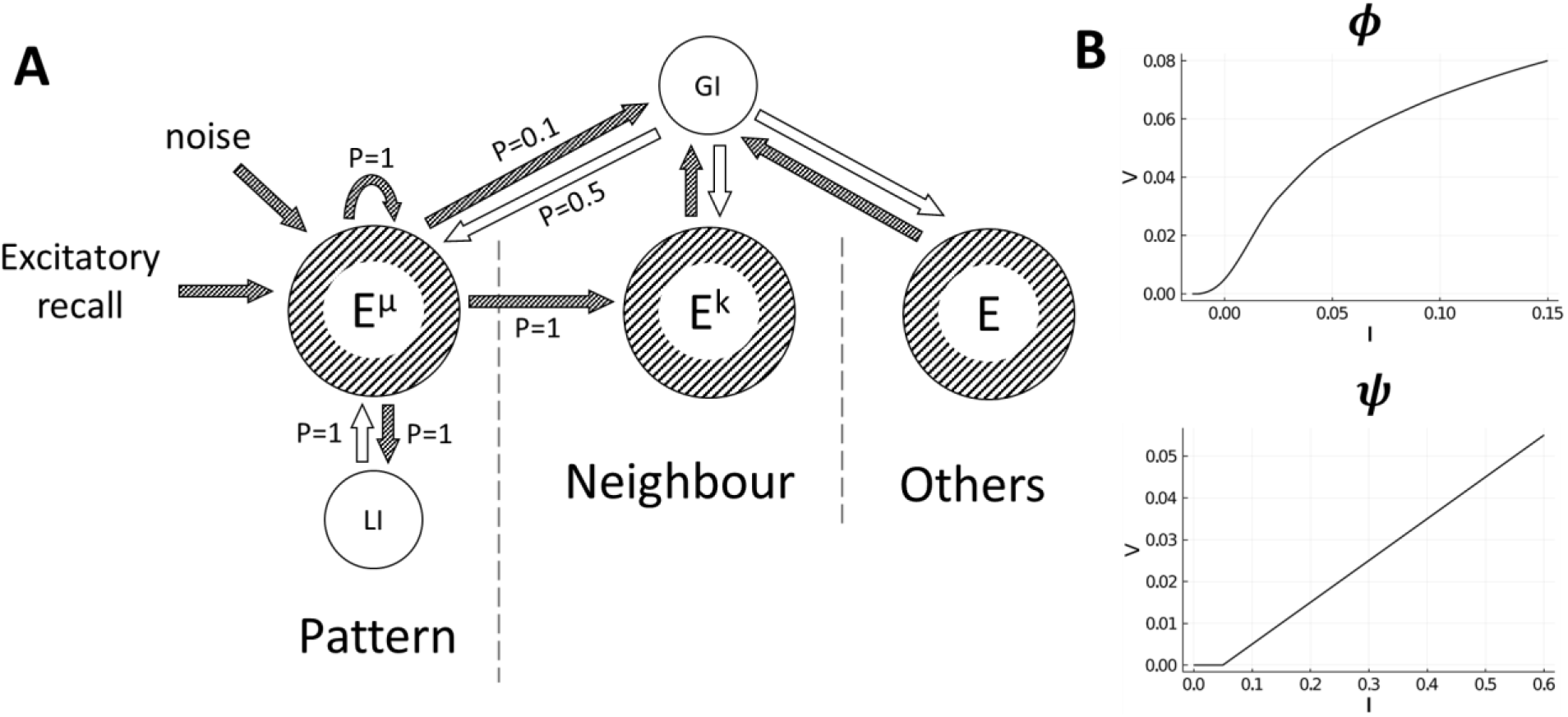
**(A)** General schematic of the model from the perspective of a single memory pattern (E^μ^) and its connections to its respective local inhibitory population (LI), neighbours (E^k^), and the global inhibitory population (GI). Connection probabilities are indicated as values of P. To retrieve a pattern, excitation is given directly to a single pattern. Gaussian noise is also applied independently to all excitatory neurons. Key: Striped/shaded arrows and circles indicate excitatory connections and populations, respectively, and unshaded arrows and circles indicate inhibitory connections and populations, respectively. **(B)** F-I curves for the excitatory neurons (*ϕ*) and inhibitory neurons (*ψ*) (from [Amit et al., 1994]). For the excitatory F-I curve, values of *I* above 0.15 are mapped to *V* = 0.08, and the inhibitory F-I curve continues linearly with the same slope for values of *I* above 0.6.

Neurons are modelled as proportions of their maximum firing rates, based on an established method [Amit et al., 1994]. At each timestep, currents are calculated for each excitatory neuron 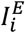, global inhibitory neuron 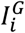, and local inhibitory neuron 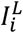

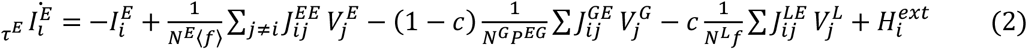

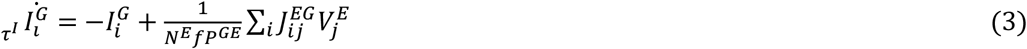

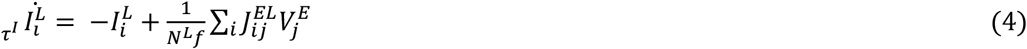

and then converted into proportions of their maximum firing rates by

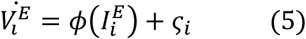

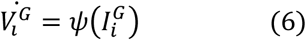

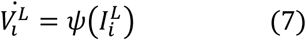

where *J*_*ij*_ is the balanced connection weight between neurons *i* and *j, τ*^*E*^ = 10*ms* and *τ*^*I*^ = 2*ms* are the time decay constants, *c* ∈ [0,1] is the local-global inhibition balance, ⟨*f*⟩ is the sum of expected firing rates based on the average degree of *M* (e.g., if *M* is a 1D chain, ⟨*f*⟩ = *f* · 1.5 = 0.015), and *P*^*EG*^ = 0.5 and *P*^*GE*^ = 0.1 are the connection probabilities from excitatory to global inhibitory neurons and global inhibitory to excitatory neurons, respectively. External input to the network is given by 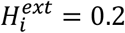, the drive given to excitatory neurons in the pattern we wish to retrieve during the stimulation window, and *ς*_*i*_ ∈ 𝒩(0, 0.0015^2^) is small Gaussian noise (independently drawn at every step, for every excitatory neuron). The F-I curves for the excitatory neurons (*ϕ*) and inhibitory neurons (*ψ*) are shown in Figure 1B and are from a previous study [Amit et al., 1994]. The network’s forward dynamics (governed by equations 2-7) are solved using the Euler method with step sizes of 0.1*ms*.

The excitatory-to-excitatory weights are considered balanced by setting 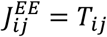. We then balance the inhibitory-to-excitatory and excitatory-to-inhibitory weights based on 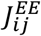. We balance the inhibitory-to-excitatory (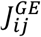 and 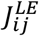) and inhibitory-to-excitatory (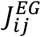 and 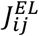) connections by calculating the sum of each excitatory neuron’s pre-synaptic input in *J*_*ij*_ and calculating the proportion of this sum compared to the mean sum of all excitatory neurons. This proportion becomes the connection weight, and obtains a mean of 1. In effect, this means excitatory neurons which receive stronger recurrent excitation than the mean excitatory neuron receive proportionally stronger local and global inhibition. Theoretically, this can be interpreted as a form of homeostatic normalisation for the purpose of excitatory-inhibitory balance.

Associative memory structures *M* = (*P, A*) with |*P*| = *p* vertices (memory patterns) and edges (memory associations), *A*, is chosen and the model is instantiated according to the above procedure. We then choose a single pattern to receive external input to all of its excitatory neurons during the stimulation window, *t* = 0*ms* to *t* = 80*ms*, after which the network is left to settle into an approximate steady-state and stopped at *t* = 500*ms* for analysis. The main variable of manipulation was the balance between local and global inhibition balance, *c*, where *c* = 0 means only global inhibition is active, *c* = 1 means only local inhibition is active, and *c* = 0.5 means there is an equal contribution of both global and local inhibition in the network.

### Analysis

To quantify the extension in the range of retrieval given by changes in *c*, we tested *M* as a 1D chain with *p* = 100. We stimulated each pattern and recorded the excitatory firing rates at *t* = 480*ms* to *t* = 500*ms*. Following an established method [Amit et al. 1994], and with *W*_*S*_ = 20*ms* being the number of timesteps being averaged, we calculate the mean 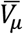 and variance 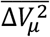 of the final firing rates for each memory *μ* by

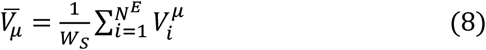

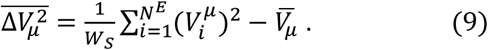

The covariance between two memories *μ* and *v* is

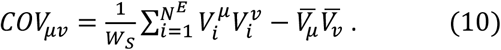

The correlation between two memories *μ* and *v* is

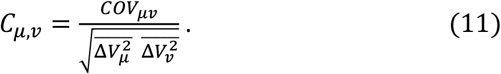

We then calculate the mean correlation between two memories at the shortest path distance *d* away from each other by

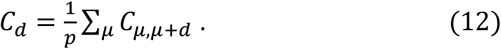

Finally, we quantify the range of retrieval 𝒟 using the following algorithm:

1. Calculate |*C*_*d*−1_ − *C*_*d*_| for all *d* > 1.
2. 𝒟 is the first value of *d* for which the next *Y* attractors have |*C*_*t*−1_ − *C*_*t*_| < *ε*. If no such 𝒟 is found, 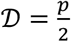.

We use *ε* = 0.05 and *Y* = 5.

We observed how the activity of the excitatory population spread through associative memory structure for different values of *c* and across time. We chose to visualise this spread in three classical graphs – Zachary’s *karate club graph* [Zachary, 1977], the *K*5*-3-chain* [Schapiro et al. 2013], and the *Tutte graph* [Tutte, 1946] – and one constructed graph representing a multi-room spatial environment which we call the *multiroom graph*. These graphs were chosen for their complexity, relation to or derivation from real-world analogues, and well-known graph theoretic features. In order to quantify the similarity between the activity of the network and graph theoretic properties in the associative memory structures, we compared the approximate steady-state activity to the community detection and classification of vertices using the label propagation algorithm [Raghavan et al., 2007]. We denote two vertices – e.g., *μ* and *v* – being members of the same community according to this algorithm with *LPA*(*v*_*μ*_, *v*_*v*_) = 1 and *LPA*(*v*_*μ*_, *v*_*v*_) = −1 otherwise. Then, the *clustering index* for a given trial and its associated associative memory structure is given by

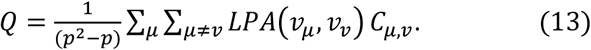

The *clustering index* is a measure of how our model’s activity corresponds to topological features of *M*. To test how the activities correspond to geometric distance for arbitrary graphs, we define a local area around a vertex in *M*. This local area is the closed *d*-neighborhood of a vertex, i.e., the set of the vertex *v* and all vertices within distance *d* as measured by their shortest path to *v*. For a choice of *d* and *v*, we construct a local area function *LA*(*μ*|*v, d*) which assigns vertices in the local area with a value of 1 and −1 otherwise. We then calculate the *geometric index* by

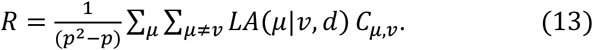

## Results

The general structure of the model is illustrated in Figure 1A. Memories are modelled as strongly-interconnected assemblies of excitatory neurons. Each memory item’s assembly is also interconnected to the assemblies of memory items which it is connected to in the associative memory structure, *M*. The associative memory structure can take on any form. Inhibition to the network is provided by two equally-sized populations: (i) a global inhibitory population, which has an excitatory to global inhibitory connection probability of 0.1 and global inhibitory to excitatory connection probability of 0.5; and (ii) local inhibitory populations (one for each excitatory assembly), which are fully connected to individual excitatory assemblies in the associative memory structure. The balance between these two activities was governed by the parameter *c*: *c* → 0 being strongly global, *c* → 1 being strongly local, and *c* = 0.5 being a balance between the two. A single trial is performed by giving a brief positive impulse (80*ms*) to a single excitatory assembly and then letting the network self-regulate its activity thereafter. This is similar to how a brief sensory stimulus of a single memory item can (even after the stimulus is removed) have persistent, representable activity and this activity can cause the retrieval of related memory items via cognition [Miyashita, 1988; MacDonald et al., 2011; Uitvlugt & Healey, 2018]. We mostly analyse the approximate steady-state reached after 500*ms*.

### Extended range of retrieval

Setting *M* as a 1D chain with *p* = 100 memory patterns, we simulated values of *c* from 0 to 1 in 0.1 steps. We found the range of retrieval extended gradually with increases to *c* (Figure 2A). At *c* = 0.7, the network showed a dramatic increase in noisy behaviour, however this slowly subsided as we increased the size of the network, indicating a finite field effect (Figure 2B). Compared to *c* = 0, which had a range of retrieval of around 5, *c* = 0.7 quadrupled this distance to 20 neighbours in distance along the 1D chain (Figure 2C). In the range of *c* > 0.7, we tested networks of sizes up to *N*^*E*^ = 128,000, *N*^*G*^ = 4,000, *N*^*L*^ = 4,000 and found that in all cases the network activity was very noisy. Due to computational limitations, we did not test larger networks, however we speculate that sufficiently large networks are likely to exhibit even greater extensions to the range of retrieval but at smaller network scales are perturbed by noise from a finite field effect.

**Figure 2.**
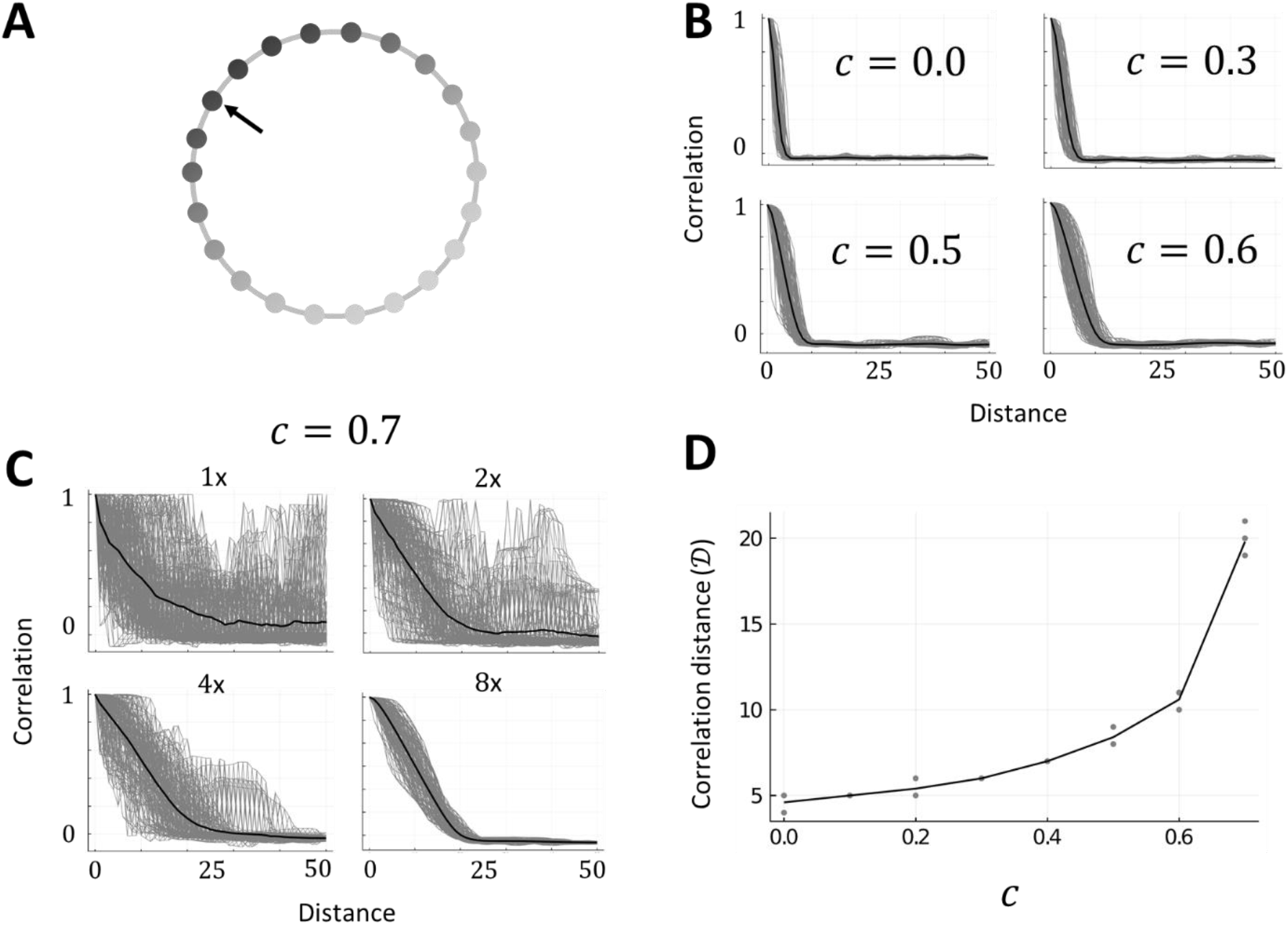
**(A)** Illustration of *M* as a 1D chain. An arrow indicates the initially stimulated memory and the shading of vertices indicates the strength of activity in each excitatory assembly (darker is more active). **(B)** Example trials showing the correlation of approximate steady-state activities of excitatory neurons with neighbouring memories in a 1D chain associative memory structure. Grey lines are single trials (n=100) and black lines are the mean of all trials. Panels show increasing the value of *c* to 0.6 (to a local inhibition dominant network configuration) approximately doubles the initial range of retrieval. **(C)** Example trials the same as (B) for *c* = 0.7 with panels showing increasing sizes of networks (starting from 1x, which is *N*^*E*^ = 4,000, *N*^*G*^ = 500, *N*^*L*^ = 500). This indicates a strong finite field effect which appears in the local inhibition dominant state. **(C)** Scatterplot showing the range of retrieval measure, 𝒟, increases with *c*. Grey dots are single trials (n=5 per value of *c*) and the black line follows the mean of trials. Trials for *c* = 0.7 were completed with *N*^*E*^ = 32,000, *N*^*G*^ = 1,000, *N*^*L*^ = 1,000, and for all other values of *c* the trials were completed with the regular network size (*N*^*E*^ = 4,000, *N*^*G*^ = 500, *N*^*L*^ = 500).

### Spread of excitation in associative memory structures

In most cases – karate club graph, *K*5-3-chain, and multiroom graph – excitation spread across a larger range of the associative memory structure when local inhibition was dominant than when global inhibition was dominant. The Tutte graph uniquely decreased the spread of excitation when activating its *central vertex* (indicated by the arrow shown in Figure 3A, third row). As in the 1D chain case, trials with values above *c* = 0.7 often had noise, although the largest graph (multiroom) had stable trials with values of up to *c* = 0.85. We also observed most graphs change in their excitatory activity most noticeably in the region of *c* = 0.5 to *c* = 0.6. We therefore chose to focus on two cases: (i) strong global inhibition (*c* = 0.1), and (ii) slightly stronger local inhibition (*c* = 0.525) (Figure 3).

**Figure 3.**
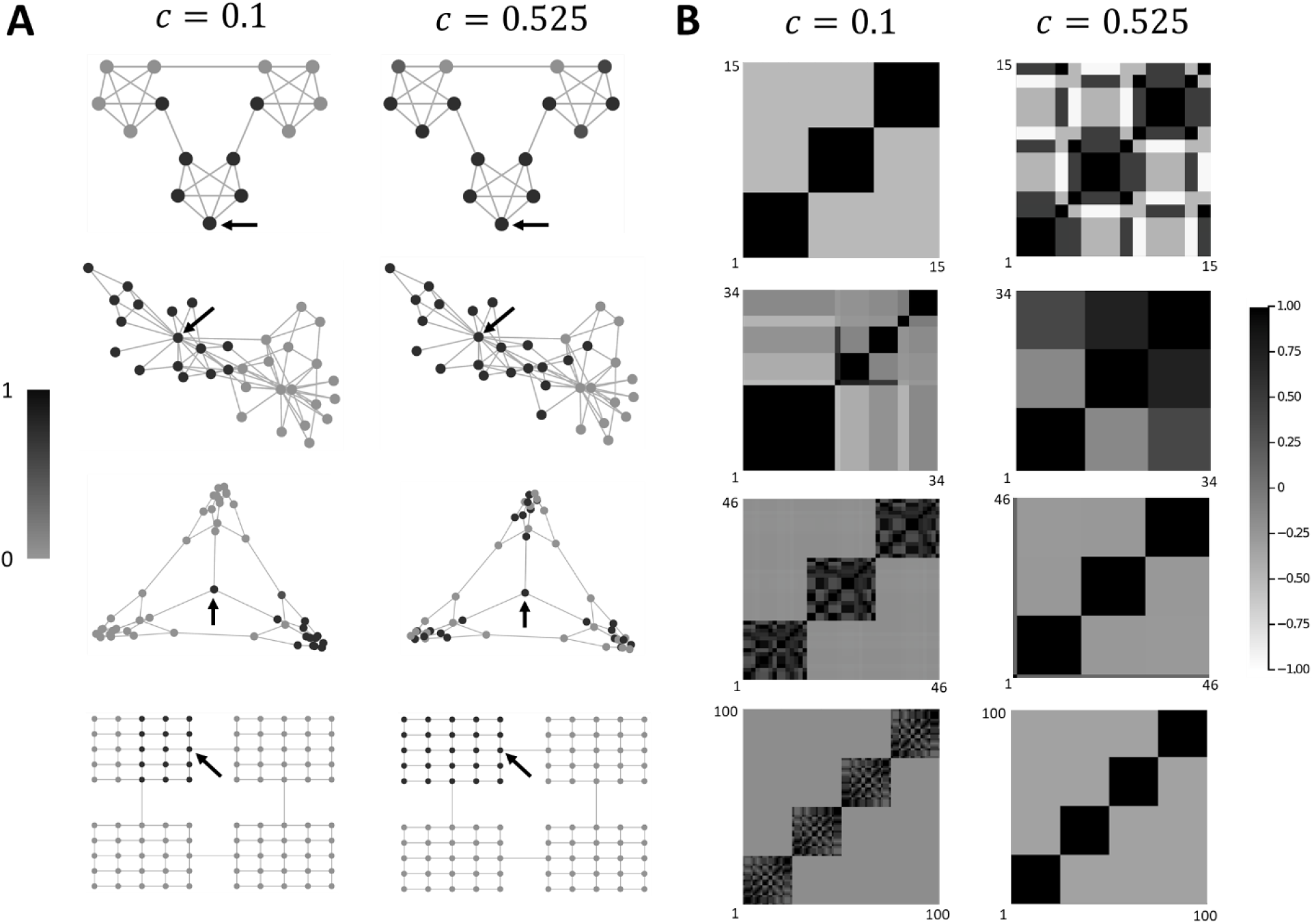
**(A)** Example trials in four different associative memory structures at two values of *c*. Vertices are shaded according to the sum of its neurons’ normalized activity (darker is more active). Arrows indicate the vertex which was stimulated at the beginning of the trial. **(B)** Correlations of approximate steady-state activities of excitatory neurons with all other vertices in the same associative memory structures and at the same two values of *c* as in panel A. Vertices have been ordered such that those with similar correlations to other vertices are adjacent to illustrate the clustering effect that naturally arises from the network’s dynamics.

**Figure 4.**
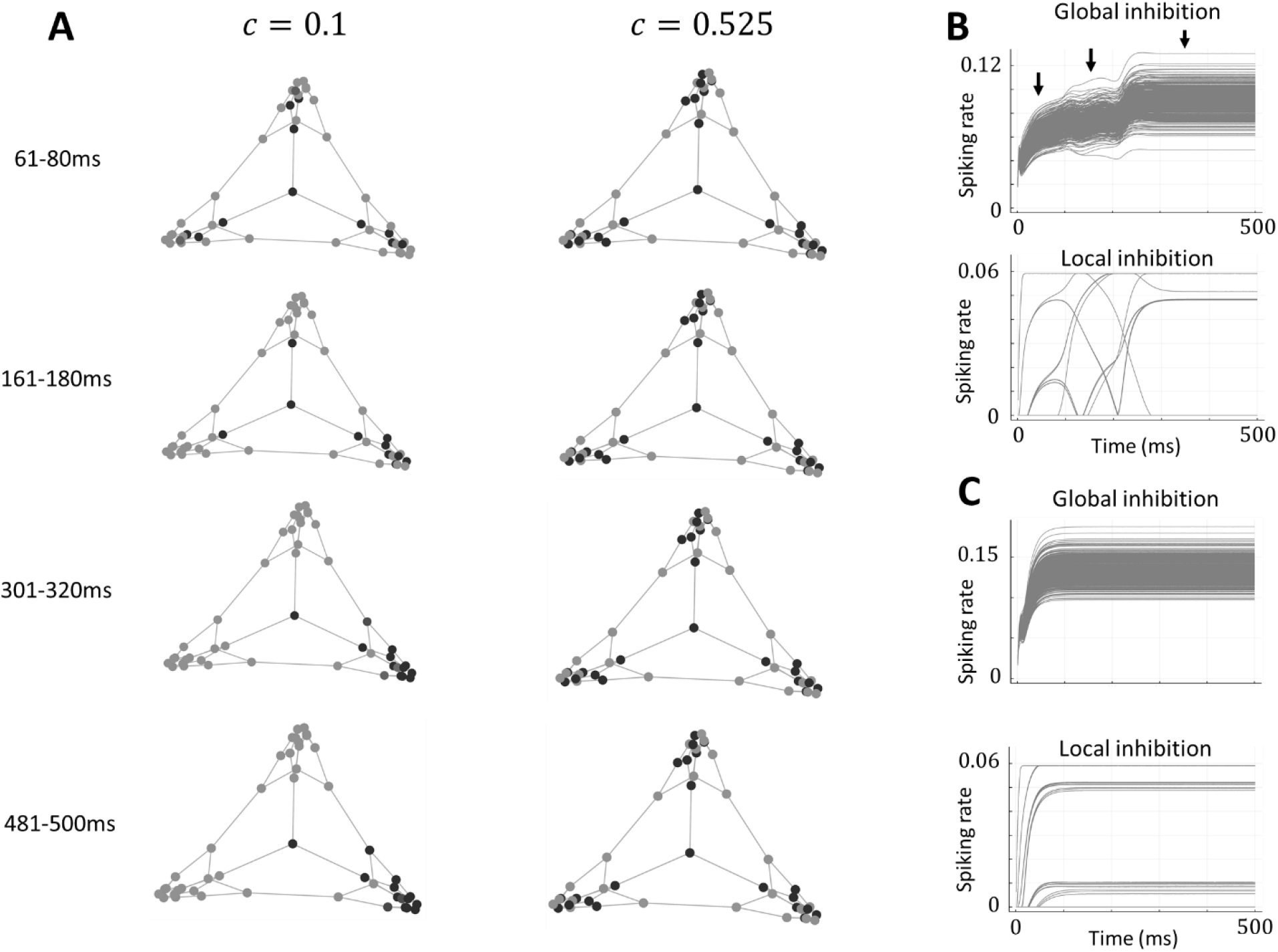
**(A)** Example trials for memory item neuron activities in the Tutte graph during different time-windows for *c* = 0.1 and *c* = 0.525. The central vertex is activated for the first 80ms of each trial. **(B)** Global and local inhibitory firing rates over time for the Tutte graph trial with *c* = 0.1 shown in (A). Arrows illustrate three distinct modes or levels of global inhibition. **(C)** Same as (B) but for *c* = 0.525.

Correlations between the vertices (assemblies) of the underlying neurons (neurons belonging to those assemblies, see eq. 11) showed different resolutions of clustering. For example, at *c* = 0.1 in the karate club graph, approximately 5 clusters of strongly correlated vertices were present, whereas at *c* = 0.525 this reduced to approximately 3 (Figure 3B, second row). Although the Tutte and multiroom graphs showed a similar trend in consolidation of clusters at *c* = 0.525, the *K*5-3-chain showed the breaking down of clusters and some strong negative correlations. We can see in Figure 3B (top row) that the graph is made up of pseudo-*K*5 subgraph – groups of 5 vertices completely connected, except for two ‘boundary’ vertices, which connect the pseudo-*K*5 subgraphs together. Within each pseudo-*K*5 subgraph, the 3 ‘core’ vertices (those which are fully connected within the pseudo-*K*5 subgraph and not the boundary vertices) remain strongly correlated with one another while the two ‘boundary’ vertices become almost equally correlated with their own pseudo- *K*5 subgraph and their neighbouring subgraph and negatively correlated with the opposite subgraph. For the well-connected core vertices, *c* = 0.525 also represents the level at which the spread of excitation almost covers the entire graph.

We also observed how excitation spreads across the associative memory structure across time, after activation of vertices of interest, in the Tutte and multiroom graphs. For the Tutte graph we chose the central vertex, which branches off into three separate ‘rooms’, and for the multiroom graph we chose a location within one of the rooms that also led through a ‘doorway’ to a neighbouring room. We chose these vertices since they represent points of behavioural interest and ecological importance in animals - they are points at which an animal may make significant choice between which room to enter, explore, or exploit. In the Tutte graph, for *c* = 0.1, there is initial activation of all three rooms (Figure 3A). This is accompanied by a general rise in global inhibition and specific increases in the activity of local inhibitory populations connected to the respective active excitatory populations. However, at this early stage, one room is slightly more dominant in overall excitation (bottom-right ‘room’ in top-left panel of Figure 3A). This dominance appears to translate into gradual and then complete activity dominance compared to the other two rooms at the later time-windows. Contrastingly, for *c* = 0.525, the activity of vertices in the Tutte graph is initially broader and this breadth of excitation is maintained steadily throughout the duration of the trial. We also see that the global and local inhibitory populations for *c* = 0.525 (Figure 3C) quickly stabilize in an approximate steady-state. In the case of for *c* = 0.1, the global inhibitory activity progresses through three distinct phases of activity (indicated by the arrows in Figure 3B): an initial rise, an unstable plateau, and finally a higher, stable plateau. Meanwhile, the local inhibitory activity for *c* = 0.1 reflects the recruitment and release of various memory items before coming to an approximate steady-state at a similar time as the global inhibitory activity.

The multiroom graph showed a similar trend in broadening and maintaining a larger range of retrieval with increases in *c*. However, possibly due to the size of the network and because the chosen vertex was located within one of the rooms (thus biasing towards activation of that room’s other vertices, unlike the central vertex in the Tutte graph), observation of the effect required an increase in *c*. For illustration of the effect, we chose *c* = 0.525 and *c* = 0.7 (Supplementary Figure 1). Interestingly, in the case of *c* = 0.7, initial broadening of the range of retrieval into the neighbouring room (through the doorway adjacent to the memory item being stimulated) was slightly reduced and the first memory pattern of the room on the opposite doorway became active later in the trial.

The clustering and geometric indices – *Q* and *R* – for each graph, at different values of *c* are given in Table 1. Since *R* depends upon a choice of distance *d* in the local area, we calculated *R* for all values of *d* (from 1 up to the diameter) and report the largest value of *R* (and its *d*). In general, the larger the value of *Q*, the more agreement between the community structure measured by label propagation and by the correlations of vertex activities in the final network states (by our model). High values of *R* indicate the final activity states are similar to geometric distance. We analyse the activity based on all neurons and a subset of neurons which reach a firing rate of at least 0.02 of the maximum firing rate during the simulation. We call this subset the *selective neurons*.

**Table 1.** Clustering and geometric indices for graphs at different values of *c*.

|  |  |  |  |  |  |  |  |  |  |
| --- | --- | --- | --- | --- | --- | --- | --- | --- | --- |
| | | Clustering indices ($Q$) | | | | Geometric indices ($R$) | | | |
|  |  | All neurons |  | Selective neurons |  | All neurons |  | Selective neurons |  |
| | Diameter | $c = 0.1$ | $c = 0.525$ | $c = 0.1$ | $c = 0.525$ | $c = 0.1$ | $c = 0.525$ | $c = 0.1$ | $c = 0.525$ |
| $K5$-3-chain | 4 | 0.404 | 0.040 | 0.643 | 0.368 | 0.337<br>($d=2$) | 0.393<br>($d=4$) | 0.557<br>($d=1$) | 0.592<br>($d=2$) |
| Karate club graph | 5 | 0.098 | -0.223 | 0.120 | -0.266 | 0.111<br>($d=2$) | 0.999<br>($d=5$) | 0.191<br>($d=4$) | 0.999<br>($d=5$) |
| Tutte graph | 8 | 0.050 | 0.270 | 0.254 | 0.244 | 0.259<br>($d=4$) | 0.389<br>($d=4$) | 0.300<br>($d=3$) | 0.464<br>($d=3$) |
| Multiroom graph | 18 | 0.115 | 0.144 | 0.095 | 0.258 | 0.210<br>($d=5$) | 0.375<br>($d=5$) | 0.210<br>($d=5$) | 0.375<br>($d=5$) |

Clustering indices (*Q*) using only the selective neurons are generally larger than for all neurons, indicating these more-active neurons generally contribute positively to clustering. This is especially noticeable when the network settles into a state where assemblies take on a wide range of values (e.g., in the *K*5-3-chain graph for *c* = 0.525). In general, the clustering indices indicate that given the size and topology of different graphs, different values of *c* have different propensities for clustering global characteristics.

Geometric indices (*R*) were generally greater than the clustering indices, indicating a greater emphasis of the geometry rather than the topology in these memory graphs at these values of *c*. Nonetheless, some topological information is captured and almost all of the geometric indices were of a comparable order as the clustering indices. In all cases, as we increased *c*, the value of the geometric index *R* strictly increased and was maximal for a strictly larger distance *d*. So, like in the 1D ring case, rebalancing the activity of inhibitory neurons such that local inhibition become more dominant extends the range of excitation.

## Discussion

Previous modelling studies have conflated excitatory and inhibitory neuron identities and learning rules [Griniasty et al., 1993; Haga & Fukai, 2019] or ignored inhibitory neurons’ functional participation [Amit et al., 1994] in associative memory structure retrieval. This work uniquely disentangles excitatory and inhibitory neurons and uses sparse excitatory assemblies to demonstrate the potential functional role of global-local inhibitory balance in a more biologically-plausible setting. In the simplistic case of a 1D memory chain (like might correspond to discrete memories in a sequence of events through time), shifting inhibition to a locally-dominant state quadrupled the range of activation or retrieval. In the case of more sophisticated memory structures, globally-dominant inhibition tended to emphasize finer scale partitions of the memory structure and consolidated strong local associations. Whereas, locally-dominant inhibition tended to capture broader scale partitions and allow excitation to extend across a larger range of the memory structure.

Stable extension in the range of retrieval appears limited due to increases in noise in strongly local-inhibition dominant states. This is likely due to a finite field effect and may indicate a necessary minimum size of local excitatory-inhibitory assemblies for such states. For example, stability in the case of *c* = 0.7 for the 1D chain case with sparsity of *f* = 0.01 required a network size four times greater than the case of *c* = 0 to maintain stability of retrieval, translating to excitatory assemblies of 160 neurons paired with 40 local inhibitory neurons. Although assemblies of approximately 300 neurons have been used in optical microstimulation experiments in sensory cortex to drive behaviour in mice [Huber et al., 2008], most recorded assemblies are on the order of tens of neurons [Harris et al., 2003; Fujisawa et al., 2008]. Among other benefits, such sparsity is accompanied by theoretical energy efficiencies [Levy & Baxter, 1996] and in associative memory models can lead to fewer spurious memories [Hoffman, 2019]. It therefore seems likely that for the described mechanism of local-global inhibition to have a stable functional effect in extending the range of retrieval, the presence of both local and global inhibition is required in finite, real-world networks with sparse assemblies.

Alternatively, it is possible this mechanism requires a hybrid sparse-dense coding schema, as has long been suggested operates in hippocampus [Barnes et al., 1990], cerebellum [Marr, 1969], and more recently in sensory areas [Laurent, 2002; Sakata & Harris, 2009]. In such a schema, sparse assemblies report their activity to densely-connected assemblies which broadcast information to other sparse assemblies. In our model, we could consider the global inhibitory population as a densely-connected assembly which broadcasts the overall level of excitation in the network to all local, sparse assemblies. It is just not excitatory, as in classic dense-sparse schemas. Through this interpretation, a reduction in the relative strength of global inhibition (as in the unstable region of *c* > 0.7) is equivalent to a gradual transition in the coding schema from sparse-dense to sparse. Thus, if the described local-global inhibition mechanism requires a sparse-dense coding schema, its instability when the coding scheme becomes sparse is expected. Associative memory structures which had more sophisticated topologies also showed unstable regions at high values of *c*, however less so when the graph was sufficiently large (such as in the multiroom graph). So, it is also possible this mechanism can be supported when the memory structure is adequately structured or large.

Extension of the range of retrieval was not simply the only apparent function of the inhibitory mechanism in sophisticated associative memory structures, the mechanism also permitted multiscale segmentation of the associative memory structure. Local-inhibition dominant states typically activated coarser topological segments of the graphs whereas global-inhibition dominant states consolidated activity in more densely associated clusters, highlighting finer topological features. These results were similar to those found in a more abstract model of binary neurons [Haga & Fukai, 2021], except that the current model was unable to eliminate the spread of excitation totally (as the more abstract model [Haga & Fukai, 2021] was capable of). This is because the current model does not include direct potentiation of excitatory weights, but rather modulation of local-global inhibitory balance. In this model, where association is embedded ubiquitously, in order to sustain highly-specific activity within a narrow range of memory items or even a single memory item, it is necessary to create very strong self-excitation within an assembly and have stronger overall inhibition with *c* → 0. This demonstrates a general limitation that in a more biologically-realistic setting it may not be possible to fully eliminate or reduce association between items embedded in an excitatory memory structure through inhibitory modulation alone. Nevertheless, such inhibitory activity may cause dissociation through plasticity and learning mechanisms – as demonstrated in numerous psychological and biological studies [Anderson, 2003; Chiu & Egner, 2015; Schmitz, 2017; Anderson & Hulbert, 2021] – which we have not investigated here.

An intriguing aspect of this inhibitory mechanism is its ability to dramatically affect not just the range of retrieval but also which parts of the memory structure become dominant given an initial stimulation. For example, it appears in global-inhibition dominant states, global inhibition drives a ‘winner-takes-all’ dynamic [Grossberg, 1973] whereby only the globally strongest memories remain active. In local-inhibition dominant states, this ‘winner-takes-all’ dynamic appears to dissipate and permit a general extension of retrieval, or a more egalitarian sharing of the winners. However, this extension can also be mediated and a ‘winner-takes-all’ dynamic can appear at the peripheries of the retrieval range, with different peripheries competing against each other (for example, see Supplementary Figure 1B). This may be considered as a global state transition from ‘winner-takes-all’ to ‘winner-shares-all’ [Fukai & Tanaka, 1997]. We therefore hypothesize an inhibitory mechanism like we have described may be used to aid in the learning or retrieval of graph-based cognitive tasks in cortical networks [Whittington et al., 2020; Wang-King et al., 2021]. Cognitive control or exploitation of this mechanism might also occur in concert with, for example, gamma oscillations, which are strongly tied to inhibitory activity [Buzsáki & Wang, 2012]. This may be especially useful when faced with competing behavioural choices and maintaining the distribution of these choices is meaningful.

Making a seemingly subtle change to the network structure by introducing some of the complexities and diversities of inhibitory neurons had a profound impact on retrieval. We have shown how this phenomenon mainly persists in a sparse, associative memory structure which obeys Dale’s law and biologically-plausible connections. We have also shown and discussed some of the potential functional roles of this mechanism in graph-based cognitive tasks and discussed how this mechanism may contribute to a type of sparse-dense coding schema.

## Supplementary material

**Supplementary Figure 1.**
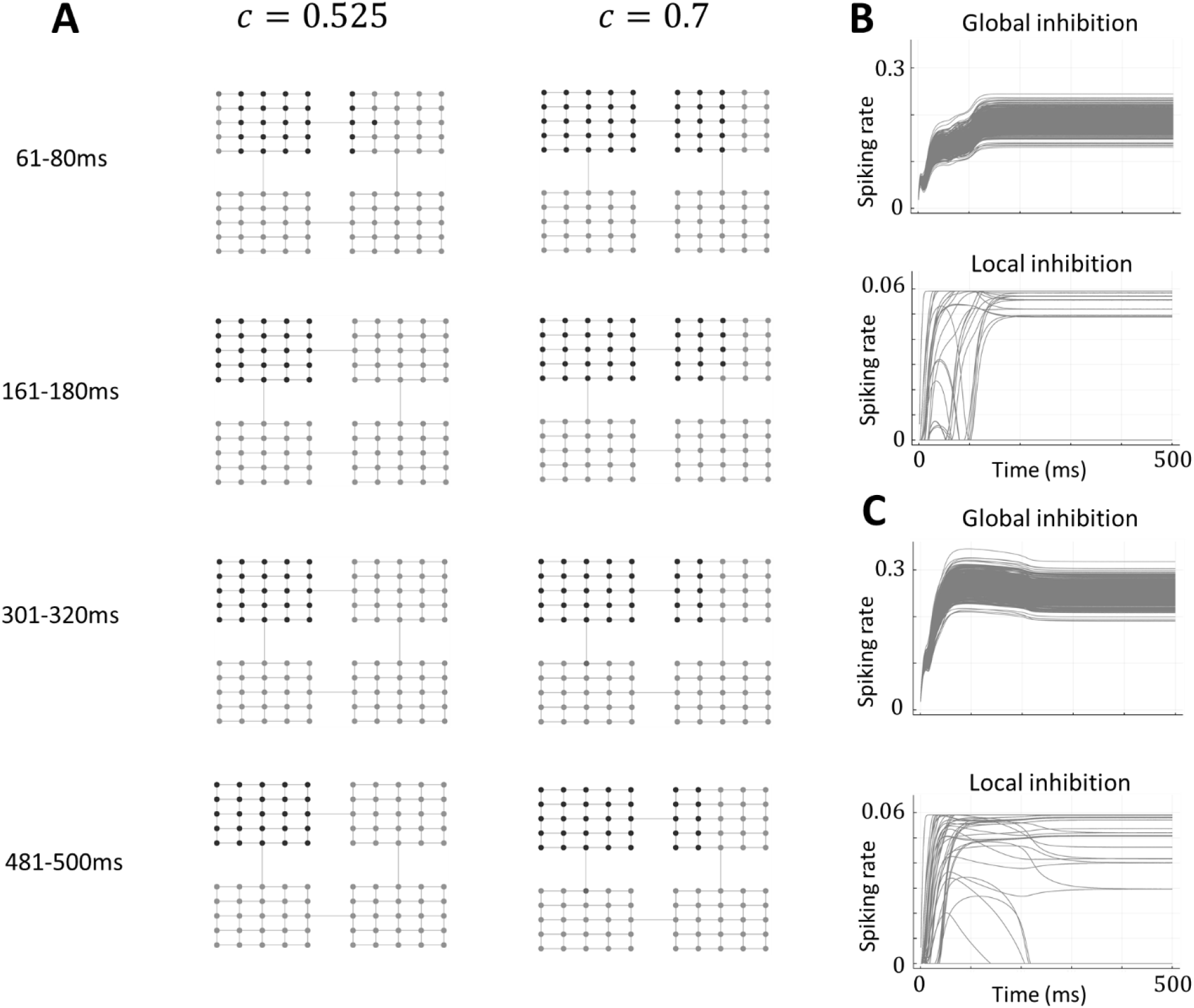
**(A)** Example trials for memory item neuron activities in the multiroom graph during different time-windows for two different values global-local inhibitory balances, *c* = 0.525 and *c* = 0.7. A vertex beside to uppermost ‘doorway’ is activated for the first 80ms of each trial. **(B)** Global and local inhibitory firing rates over time for the multiroom graph trial with *c* = 0.525 shown in (A). **(C)** Same as (B) but for *c* = 0.7.

## Notes

### Competing Interest Statement

The authors have declared no competing interest.

